# Zebrafish do not have calprotectin

**DOI:** 10.1101/2024.06.25.600640

**Authors:** Kona N. Orlandi, Michael J. Harms

## Abstract

The protein heterodimer calprotectin and its component proteins, S100A8 and S100A9, play important antibacterial and proinflammatory roles in the mammalian innate immune response. Gaining mechanistic insights into the regulation and biological function of calprotectin will help facilitate patient diagnostics and therapy and further our understanding of the host-microbe interface. Recent literature has identified zebrafish s100a10b as zebrafish calprotectin based on sequence similarity, genomic context, and transcriptional upregulation during the immune response to bacterial infections. The field would benefit from expanding the breadth of calprotectin studies into a zebrafish innate immunity model. Here, we carefully evaluated the possibility that zebrafish possess a calprotectin. We found that zebrafish do not possess an ortholog of mammalian S100A8 or S100A9. We then identified four zebrafish s100 proteins— including s100a10b—that are expressed in immune cells and upregulated during the immune response. We recombinantly expressed and purified these proteins and measured the antimicrobial and proinflammatory characteristics. We found that none of the zebrafish proteins exhibited activity comparable to mammalian calprotectin. Our work demonstrates conclusively that zebrafish have no ortholog of calprotectin, and the most plausible candidate proteins have not convergently evolved similar functions.

## INTRODUCTION

Calprotectin plays critical roles in mammalian innate immunity (Wang et al. 2018). Upon release from immune cells after damage or during an immune response, calprotectin exerts antimicrobial activity by sequestering transition metals essential for microbial growth in the extracellular matrix, also known as nutritional immunity (Urban et al. 2009; Damo et al. 2013; Hayden et al. 2013; Brophy, Hayden, and Nolan 2012; Gagnon et al. 2015; Nakashige et al. 2015; Clark et al. 2016; Baker et al. 2017; Hadley, Gu, and Nolan 2018; Besold et al. 2018). The calprotectin protein exists as both a heterodimer and heterotetramer complex formed by two calcium binding proteins, S100A8 and S100A9, which also form homodimers to a low extent.

Extracellular S100A8 and S100A9 homodimers, as well as the heterodimer calprotectin, can amplify the inflammatory response by activating Toll-like receptor 4 (TLR4) and the Receptor for Advanced Glycation End-products (RAGE), promoting cytokine expression and immune cell migration (B. Chen et al. 2015). S100A8 and S100A9 also perform biological functions distinct from calprotectin (Roth et al. 1993; Goebeler et al. 1995; Vogl et al. 2004; B. Chen et al. 2015; Lackmann et al. 1992; Ryckman et al. 2003; Passey et al. 1999; Manitz et al. 2003). Because of its high concentration and many roles in immune function, calprotectin has become a well-validated, non-invasive biomarker of inflammation (Jukic et al. 2021).

Given the importance of calprotectin, there is interest in developing new models to study its function. One attractive model is the zebrafish, which is increasingly being used to understand the molecular mechanisms of immune functions (Trede et al. 2004; Levraud, Rawls, and Clatworthy 2022). As vertebrates, zebrafish share much of their physiology and molecular components with humans. They also have exceptional experimental advantages: well-established genetic tools, optically transparent larvae (making it possible to visualize tagged molecules and microbes in real-time in live fish), and rapid generation times (Grunwald and Eisen 2002). They are particularly useful for studying innate immunity because they rely solely on innate immune responses until 4-6 weeks post-fertilization, when their adaptive immune system is morphologically and functionally mature (Willett et al. 1999; Davidson and Zon 2004; Trede et al. 2004; Lieschke and Currie 2007).

Recently, the zebrafish protein s100a10b was identified as “calprotectin” (Farr et al. 2022; Nag et al. 2022). This identification was based on genomic context and sequence similarity to human S100A8. Based on transcriptional analyses, it was proposed that this protein plays a role in the zebrafish response to *Vibrio cholerae* and *Escherichia coli* infections (Farr et al. 2022; Nag et al. 2022). Further, there are commercial ELISA antibodies marketed as targeting “fish calprotectin,” implying that this protein can be found in zebrafish.

Despite the power of the zebrafish model system, it can be challenging to map zebrafish biology to human biology. Over 400 million years of evolution have allowed the divergence, emergence, and loss of proteins and protein functions between these species, often making the comparison difficult. One of the most important considerations is whether the genes being compared between species are, in fact, the same genes. Are they the result of speciation (orthologs) which often have very similar functions, or did they arise by gene duplication (paralogs) which often have very different functions?

Establishing gene orthology is particularly challenging for S100 proteins, as they form the largest subgroup within the superfamily of proteins carrying the Ca^2+^-binding EF-hand motif. Humans have 24 S100 genes (H. Chen et al. 2014; Marenholz, Heizmann, and Fritz 2004); zebrafish have 14 (Kraemer, Saraiva, and Korsching 2008; Zhang et al. 2019). Many of these S100 genes are in dense blocks of tandem repeats. There is no annotated S100A8 or S100A9 ortholog in the zebrafish genome; however, there are several zebrafish s100 genes in a similar genomic location to that of human S100A8 and S100A9.

We set out to find phylogenetic, biochemical, or biological evidence of calprotectin (or calprotectin-like) activity in zebrafish s100 proteins. Through a careful review of existing phylogenetic literature, we confirm that fish do not have a calprotectin ortholog: both S100A8 and S100A9 evolved in mammals 250 million years after the divergence of tetrapods and ray-finned fishes. We support this phylogenetic result through a comparative synteny analysis of S100 genes in zebrafish and human genomes. We also investigate the possibility that fish convergently evolved a calprotectin-like s100 protein. We used published single-cell RNAseq data to identify zebrafish s100 proteins expressed in immune cells. We then recombinantly expressed and purified four of these proteins—including zebrafish s100a10b, the protein previously identified as fish calprotectin in the literature—and experimentally tested their antimicrobial and pro-inflammatory activities. None of the proteins give measurable activity.

We conclude that zebrafish have neither a vertically inherited ortholog of calprotectin, nor an obvious candidate protein that convergently evolved similar function. Our results highlight the hazard of relying solely on sequence similarity and genomic placement to identify genes, and demonstrate the importance of an explicitly evolutionary lens with careful functional analyses when mapping results from model organisms to human biology.

## RESULTS

### Zebrafish s100s are distantly related to human S100A8 and S100A9

We started by looking for phylogenetic evidence that fish have a protein orthologous to mammalian S100A8 or S100A9. Orthologous proteins arise by speciation and are thus the same gene in the species being compared. Paralogous proteins arise by gene duplication and often exhibit gain or loss of function from the ancestral state (Ohno 1970), establishing themselves as new proteins. **Fig 1A** summarizes the evolutionary history of S100s. This tree was built referencing several published phylogenetic analyses of the family, including two from our group (Kraemer, Saraiva, and Korsching 2008; Wheeler et al. 2016; Loes, Bridgham, and Harms 2018; Zimmer et al. 2013). The phylogeny at the top shows the current best estimate of the S100 gene tree; the phylogeny on the left shows the evolutionary history of bony vertebrates. Each circle denotes the S100 gene observed in at least one member of the taxonomic group on the left.

This evolutionary tree indicates that S100A8 and S100A9 evolved by gene duplication from a single gene in the ancestor of amniotes. Reptiles and birds preserve a single protein (MRP-126), while mammals expanded it into three proteins (S100A8, S100A9 and S100A12). The closest evolutionary relatives of these proteins are S100A7, S100A7A and S100A15. Like S100A8, S100A9 and S100A12, these arose by duplication of a single gene in the ancestor of amniotes. The reptile/bird protein MRP-126 is the earliest diverging protein known to exhibit nutritional immunity and/or Toll-like receptor 4 activation in functional assays (Bozzi and Nolan 2020; Loes, Bridgham, and Harms 2018). These observations indicate that calprotectin evolved in amniotes ∼320 million years ago.

In contrast, zebrafish s100a10b (the putative zebrafish calprotectin) falls into a clade with amniote proteins S100A10 and S100A11. This is one of the earliest S100 protein subfamilies to evolve, with orthologs present in species ranging from tetrapods to jawless fishes. This group of S100 proteins thus diverged from the lineage that led to mammalian S100A8 and S100A9 at least 563 million years ago, in the last common ancestor of humans and lampreys. Further, after this speciation event, there were at least two more gene duplications on the lineage leading to S100A8 and S100A9. S100a10b is therefore a different gene than S100A8 or S100A9.

**Fig 1:**
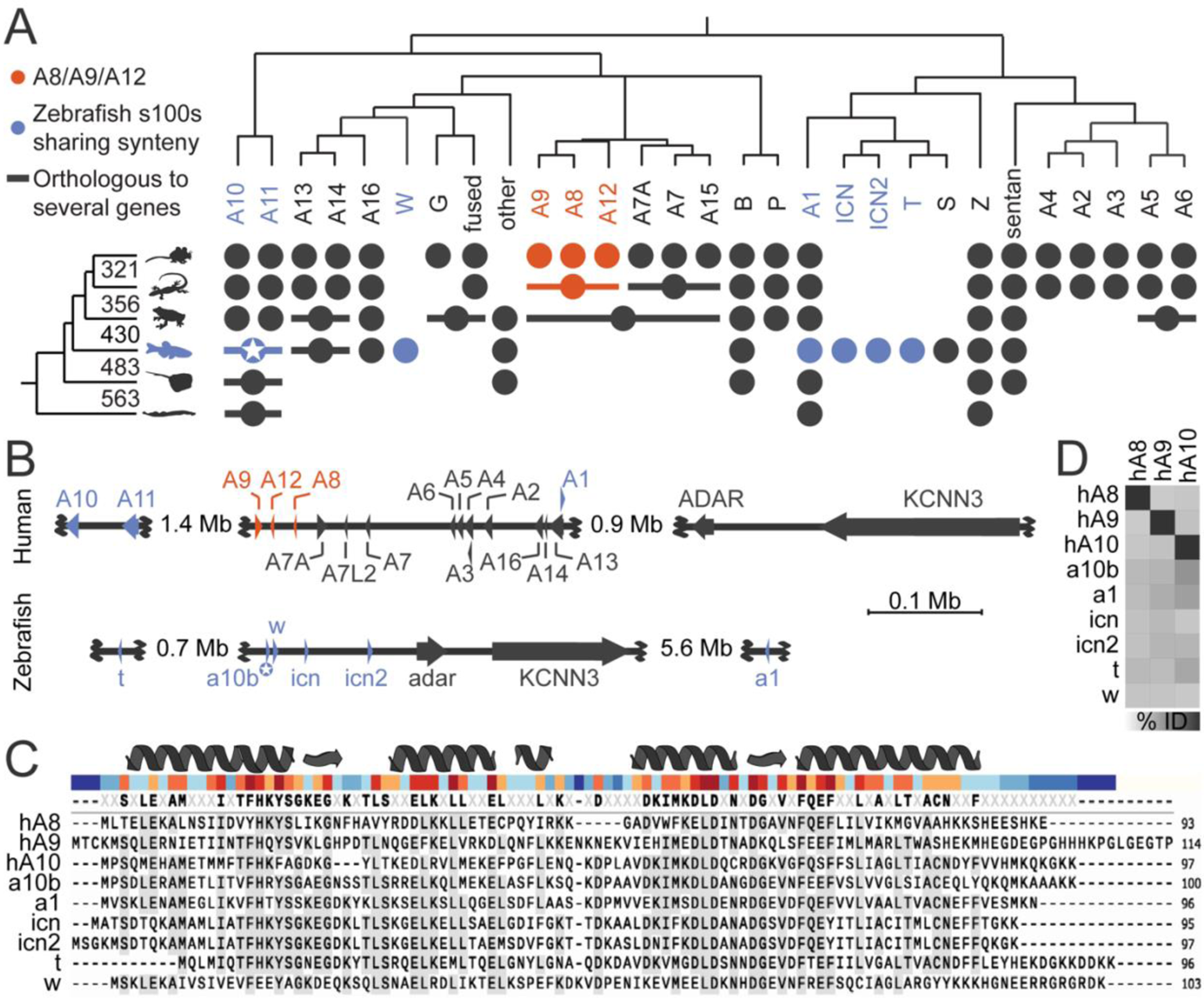
Phylogenetic analyses reveal there is no calprotectin ortholog outside of amniotes. (A) S100 gene tree adapted from Wheeler et al., 2016 shows the evolutionary relationships determined for S100s across vertebrates. The phylogeny on the left shows the relationships between species, with branch point times noted in millions of years ago; the phylogeny on the top shows the estimated S100 gene tree. Circles denote S100 genes from the phylogeny at the top found in at least one member of the taxonomic group from the left. A horizontal line through a circle indicates a single gene that is co-orthologous to multiple S100 genes found in mammals. S100A8, S100A9, and S100A12 form a clade specific to amniotes (orange). Zebrafish genes in the syntenic region shown in panel B are shown in blue. Zebrafish s100a10b, which has been treated as a calprotectin ortholog, is denoted with a star. (B) Syntenic regions of human chromosome 1:151,982,915-154,870,281 (top) and zebrafish chromosome 16:22,682,280-29,387,215 (bottom) identified by ENSEMBL. Arrows denote relative gene length and orientation. Human S100A8, S100A9, and S100A12 are shown in orange; zebrafish s100s and their human orthologs are shown in blue. The non-S100 genes adar/ADAR and KCNN3 are diagnostic for the syntenic region. Not all genes in the region are depicted. (C) A multiple sequence alignment of human S100A8, S100A9 and S100A10 compared to the zebrafish s100s from a similar genomic context. At the top, secondary structure features of human S100A8 are shown. Under this, the bar is colored by amino acid conservation from the alignment (blue=low, red=high). The consensus sequence from all sequences in the alignment is shown above the individual protein sequences. Amino acids found in a particular position in at least 50% of the sequences shown are shaded. The antigen for the “Fish Calprotectin” antibody was raised against the peptide boxed in yellow. (D) A pairwise percent identity matrix of S100 protein sequences from C (darker box indicates higher identity). S100 pair identity values for zebrafish s100s compared to human S100A8 range from 29.4-34.4%, where both s100a10b and s100t score 34.4%. Values comparing zebrafish proteins to S100A9 range from 29.1-39.6% with the highest identity shared with s100a1.

### Chromosome placement indicates a shared origin but complicated evolution of S100s in humans and zebrafish

To cross-validate the lack of evidence for vertical inheritance from published phylogenies, we used syntenic analysis to identify zebrafish s100 genes in a similar genomic location to human S100A8 and S100A9. We used ENSEMBL to identify the zebrafish genomic region most similar to human chromosome Chr 1:152-155M, which encodes 19 of the 24 human S100 proteins, including S100A8 and S100A9. This region corresponded to zebrafish chromosome 16 (S1 Fig). Specifically, human Chr 1:154.6M-154.7M and zebrafish Chr 16:23.5M-23.7M cover the KCNN3 and ADAR genes adjacent to tandem repeats of S100 genes in both species (**Fig 1B**).

The existence of this shared cluster indicates that a handful of S100 genes were in this genomic context at least in the bony vertebrate ancestor ∼430 million years ago, as established in previous work (Kraemer, Saraiva, and Korsching 2008). The syntenic relationships, however, also give evidence for extensive evolution after the divergence of bony fishes and tetrapods: the orientation and placement of genes are different, and several orthologs to human S100s are missing from this genomic location but present on other zebrafish chromosomes. Further, most of the zebrafish s100s in this region appear to be teleost-specific duplicates (Kraemer, Saraiva, and Korsching 2008). This includes ictacalcin (icn), icn2, s100t, s100s, and s100w. The only clear orthologs to human proteins are zebrafish s100a10b and s100a1 (**Fig 1A-B**).

### Human calprotectin and zebrafish s100 protein sequences share low sequence identity

Previous workers identified zebrafish s100a10b as calprotectin using human S100A8 as a query in a BLASTP search against the zebrafish proteome (Nag et al. 2022). In line with their results, we found that using human S100A8 as a BLASTP query against the NCBI RefSeq database (v. 224) yielded s100a10b as the top zebrafish hit (e-value: 8E-17, percent identity: 36.4%). The zebrafish icn2 and icn proteins were the next most similar hits. We assessed the quality of the hit by reciprocal BLAST, meaning we used the zebrafish s100a10b protein sequence as a query against the human proteome on NCBI. This yielded human S100A1 (9E-34; 55.9%) as the top hit, not S100A8. In fact, S100A8 (5E-16; 36.4%) and S100A9 (4E-18; 42.5%) were far down the hit list, following human proteins S100A10, S100Z, S100P, S100A11, S100B, S100A4, S100A12, S100A5, S100A6, and S100A2. This is consistent with previous phylogenetic analyses that place S100A8 and S100A9 as relatively distant paralogs to zebrafish s100a10b (Fig 1A).

To evaluate sequence similarity and identity, we aligned zebrafish s100 protein sequences from the syntenic region to human S100A8, S100A9, and S100A10 sequences (**Fig 1C**). As expected, there is high conservation at sites that form the EF-hand and pseudo-EF-hand calcium-binding domains of the S100 proteins. There is low conservation in the region connecting the two EF-hands and at the termini. We next determined the sequence identity shared between zebrafish s100s and human S100A8, S100A9, and S100A10 (**Fig 1D**; S1 Table). We find similarly low levels of shared identity between human S100A8 and S100A9 and the zebrafish s100s. Overall, there is no obvious candidate zebrafish s100 that is like calprotectin by sequence similarity or identity.

The sequence alignment also allowed us to ask what zebrafish s100 protein(s) might be recognized by the commercially available “Fish Calprotectin” ELISA Kit from MyBioSource. This kit was made with antibodies raised against a 20 amino acid partial peptide of a human calprotectin-like protein (GenBank: AAB33355.1), which forms the N-terminal helix and beginning of the EF-hand 1 domain of human S100A8 (**Fig 1C**, yellow box). Eight to nine residues in this helix are highly conserved in several zebrafish s100s including s100b, s100a10b, s100a10a, s100a1, s100z, s100s and s100w as well as several other unrelated proteins. This suggests the antibody may have broad, non-specific interactions with multiple zebrafish proteins.

### Single cell RNA sequencing dataset analyses point to candidate zebrafish s100 proteins upregulated in immune cells during an immune response

Our bioinformatic analyses revealed that no zebrafish s100 protein is orthologous to human calprotectin; however, it is possible that an s100 protein convergently evolved calprotectin-like activity. To investigate this possibility, we identified zebrafish s100s that share a similar expression profile to calprotectin. Calprotectin is expressed constitutively in mammalian neutrophils, monocytes, and several epithelial cell types and is upregulated upon infection and injury (Edgeworth et al. 1991; Odink et al. 1987). We queried existing zebrafish single cell RNA sequencing (scRNAseq) datasets for zebrafish s100s expressed in immune cells and upregulated in response to injury.

We used the UCSC cell browser to visualize a zebrafish development dataset (NCBI Bioproject: 564810) (Farnsworth, Saunders, and Miller 2020; Speir et al. 2021) and assessed constitutive immune cell expression in whole fish 1-, 2-, and 5-days post-fertilization. Of the genes that share genomic context with S100A9 (Fig 1B), we found that s100a10b, icn, icn2, s100w and s100a1 are expressed in immune cells of developing zebrafish (**Table 1**; S2 Fig). S100t, also in this genomic region, shows very low expression in immune cells. We found that five s100 genes from other zebrafish chromosome locations show some expression in immune cells: s100v1, s100v2, s100u, s100z, and s100a11. Finally, the remaining annotated zebrafish s100 genes—s100s, s100b, and s100a10a—appear in very few cells within these clusters.

**Table 1:** Single-cell RNAseq profiles for zebrafish s100s syntenic to human calprotectin.

| Gene name | Linkage Group and Syteny | Ortholog Call | Developmental dataset | Injury dataset |
| --- | --- | --- | --- | --- |
| **s100a1** | Chr 16<br>~syntenic | Z, A1 | Immune cells | Widely upregulated, low-level expression, hematopoietic cells |
| **s100a10b** | Chr 16<br>syntenic | A10 | Immune cells | Widely upregulated, high expression, hematopoietic cells |
| **s100w** | Chr 16<br>syntenic | A13, A14, A16 | Immune cells | Widely upregulated, moderate expression, hematopoietic cells |
| **icn** | Chr 16<br>syntenic | A1 | Immune cells | Widely upregulated, high expression, hematopoietic cells |
| **icn2** | Chr 16<br>syntenic | A1 | Immune cells | Widely upregulated, high expression, hematopoietic cells |
| **s100t** | Chr 16<br>~syntenic | A1 | Low in immune cells | Epithelial cells |

To evaluate whether these proteins are expressed during the innate immune response to injury, we used a fin regeneration scRNAseq dataset (NCBI GEO accession number GSE137971) (Hou et al. 2020). In this dataset, cells were isolated from regenerating adult zebrafish caudal fins at 1-, 2-, and 4-days post-amputation. The macrophage marker *mpeg1.1* is highly expressed in hematopoietic cells during the response to fin clip injury and is also expressed in other cell types. The neutrophil marker *mpx* was only detected at very low levels in four basal epithelial cells. We see that s100a10b, icn, icn2, s100w, and s100a1 are expressed in hematopoietic cell clusters, albeit s100a1 to a lesser degree (**Table 1**; S3 Fig). S100t only appears twice in the hematopoietic cells sequenced but shows wider expression in epithelial cells. Zebrafish s100s from other regions of the zebrafish genome show varying levels of expression in hematopoietic cells after injury. Notably, all s100 genes from outside of zebrafish chromosome 16 were expressed to a lesser degree than s100a10b, icn, icn2 and s100w. From these scRNAseq dataset analyses, we determined that zebrafish s100a1, s100a10b, s100w, icn and icn2 showed the highest gene expression profile similarity to calprotectin.

### Recombinant zebrafish s100 proteins fold and interact with calcium

We chose to functionally characterize the zebrafish s100 proteins which seemed most promising to behave like calprotectin based on genomic context and gene expression profiles: s100a10b, s100a1, s100w and icn. We left out icn2 because it is very similar to icn by all metrics including sequence identity (87.37%: they differ by 14 amino acids, 8 of these at the termini; Fig 1). The protein sequences and plasmids used for recombinant expression are shown in S2 Table.

We started by structurally characterizing the four selected zebrafish proteins. We used AlphaFold2 to predict structures for all four proteins (Jumper et al. 2021; Evans et al. 2021; Mirdita et al. 2022). Overlaying the predicted structures with the crystal structure of human calprotectin (RCSB ID 5W1F) shows high predicted structural similarity (**Fig 2A**). High α-helical content is a shared feature of all known S100 proteins, as well as the predicted zebrafish s100 structures. We tested whether this held for our selected zebrafish s100 proteins recombinantly expressed and purified from *E. coli.* We measured their secondary structure content by far-UV circular dichroism (CD). This revealed signal minima at 208 and 222 nm consistent with primarily α helical structures (**Fig 2B**).

**Fig 2:**
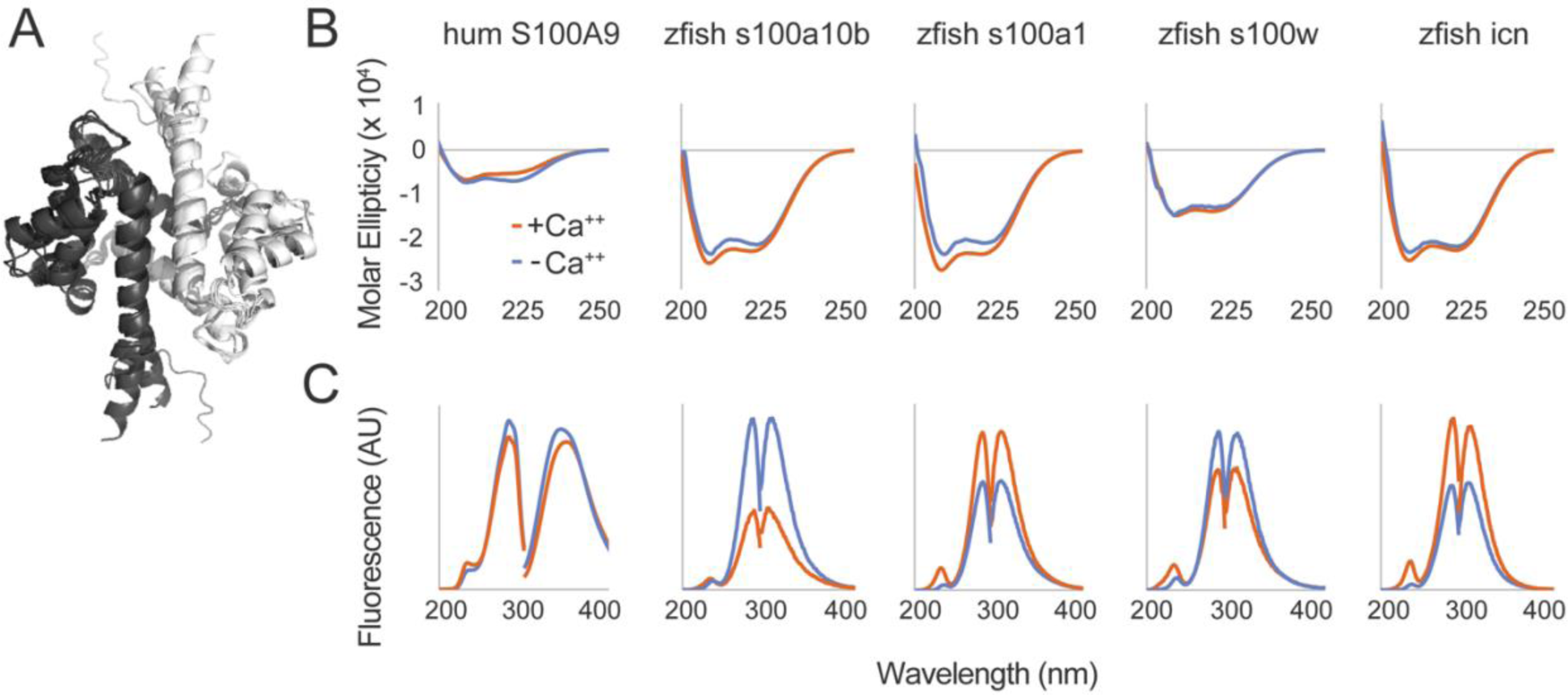
Recombinantly expressed zebrafish s100 proteins are folded and respond to calcium. (A) Overlaid AlphaFold2 structure predictions for all zebrafish s100 homodimers in this dataset, as well as the human S100A8/S100A9 heterodimer (5W1F RCSB ID). Different chains of each homodimer are shown in black or white. (B) Far UV circular dichroism spectra for each protein in the presence of 2 mM Ca^++^ (orange) and then adding 5mM EDTA (blue). Units are in molar ellipticity (deg×cm^2^/dmol) over wavelength (nm). (C) Fluorescence excitation and emission spectra for each protein in the presence or absence of calcium (orange and blue, respectively). The fluorescence units are arbitrary; the x-axis is wavelength in nanometers. Excitation spectra were collected while observing fluorescence at the maximum emission wavelength; emission spectra were collected while exciting at the maximum excitation wavelength.

Most S100 proteins also bind calcium and undergo a conformational change exposing a hydrophobic binding surface (Bhattacharya and Chazin 2003; Santamaria-Kisiel, Rintala-Dempsey, and Shaw 2006). We tested whether this held for the four zebrafish s100 proteins by measuring calcium-induced changes in protein secondary and tertiary structure by far-UV CD and intrinsic fluorescence, respectively. We found that all four recombinantly expressed zebrafish proteins exhibited evidence of calcium-induced conformational change (**Fig 2B-C**).

Upon addition of saturating calcium (orange), zebrafish s100a10b and s100a1 exhibited an increase in helical content, while s100w and icn, in contrast, show little change in secondary structure (**Fig 2B**). The intrinsic fluorescence of all four proteins, however, responded to calcium (**Fig 2C**). Intrinsic fluorescence captures changes in the local chemical environments of tyrosine and tryptophan residues, suggesting that calcium binding induces a change in the tertiary structure of all measured s100 proteins. This is consistent with the canonical calcium-induced rotation of the third helix relative to the other helices of S100s (Bhattacharya and Chazin 2003).

Taken together, these results show that these four zebrafish s100 proteins are folded, bind to calcium, and undergo the calcium-induced conformational changes expected for members of the family.

### Zebrafish s100s do not exhibit nutritional immunity characteristics like human calprotectin

One of the most important biological functions of human calprotectin is antimicrobial activity via nutritional immunity. We evaluated the antimicrobial abilities of each of the four zebrafish s100 proteins against human-derived *Stapholococcus epidermidis* and zebrafish-derived *Vibrio ZWU0020* and *Aeromonas ZOR001* strains. *S. epidermidis* was previously shown to be susceptible to human calprotectin (Harman et al. 2020; Hadley, Gu, and Nolan 2018); the response of the zebrafish-derived strains is unknown. **Fig 3A** shows the dose-dependent antimicrobial activity of human calprotectin against each strain over 13 hours in nutrient rich media across three biological replicates. For all three strains, increasing amounts of calprotectin (from blue to green) leads to decreased final OD_600_ values (indicated by black arrows). We quantified this response by measuring the difference in area under the OD_600_ curve from 0-13 hrs with and without calprotectin (ΔAUC). We then plotted ΔAUC as a function of protein concentration. A negative ΔAUC value indicates growth inhibition at the indicated s100 concentration, while a zero or positive value indicates no antimicrobial activity. This revealed a calprotectin-dependent decrease in growth for all three bacterial strains (**Fig 3B**, yellow curves).

**Fig 3:**
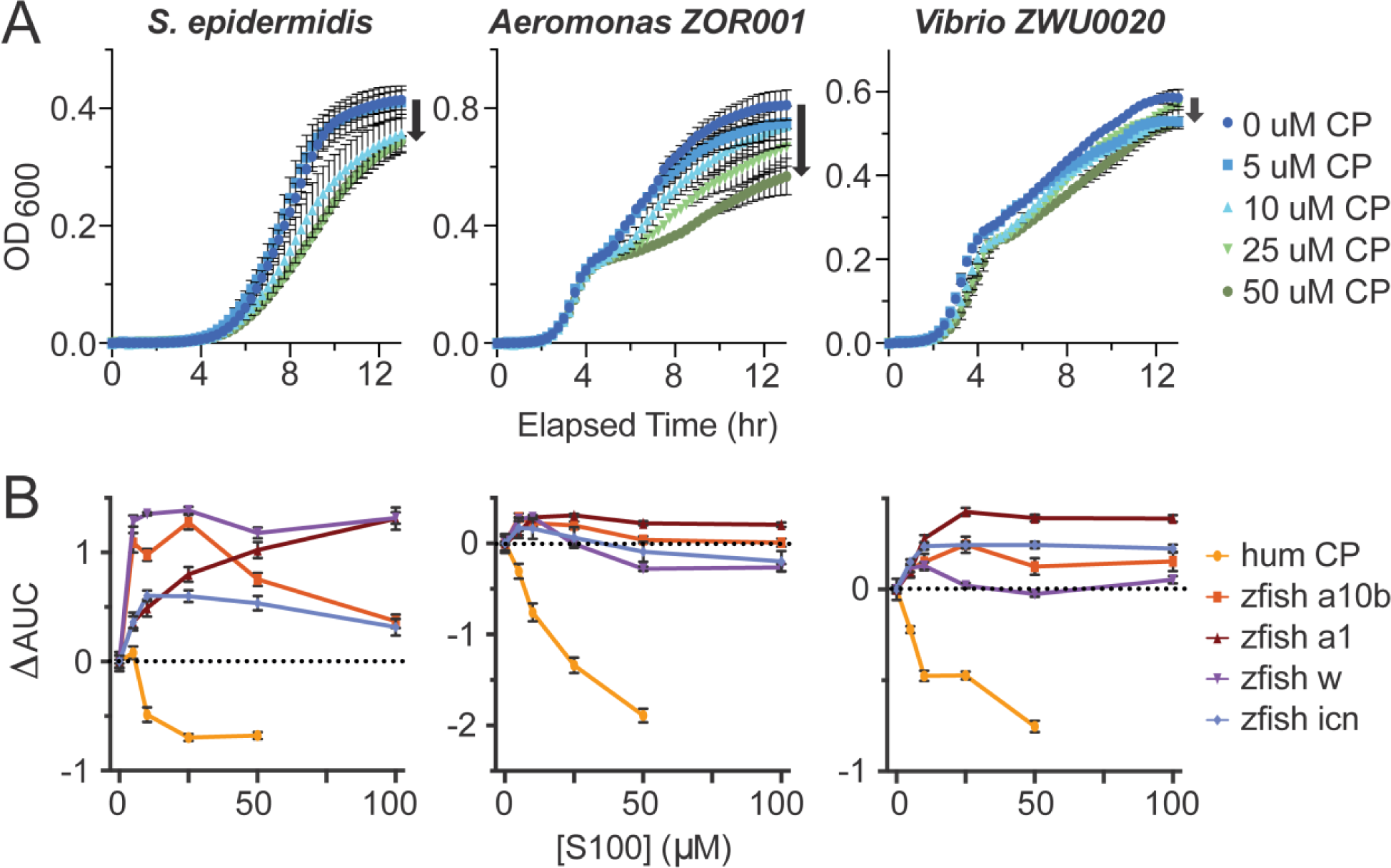
Zebrafish s100s do not exhibit nutritional immunity activity like human calprotectin. (A) Dose dependence of human calprotectin challenge on human and zebrafish commensal bacteria. Each column specifies which bacteria was used for the set of nutritional immunity assays: a human-derived Gram-positive *Stapholococcus epidermidis* and two zebrafish-derived Gram-negative bacteria, *Aeromonas* strain: *ZOR001* and *Vibrio* strain: *ZWU0020.* Bacterial growth was measured by OD_600_ over 13 hours with challenge by increasing doses of human calprotectin noted in the legend on the right. Concentration increases from dark blue to dark green and black arrows indicate how growth is affected as calprotectin increases. Error bars indicate standard error of three biological replicates. (B) Zebrafish s100 dose effects on human and zebrafish commensal bacterial growth compared to human calprotectin. Each datapoint shows the change in the area under the curve from the absence of s100 protein to the indicated s100 concentration, measured from growth curves like those shown in panel A. The dotted line at zero represents no effect on bacterial growth.

To assess the nutritional immunity capacity of the zebrafish s100s, we performed identical experiments using each of the four proteins (OD_600_ dose response curves for zebrafish s100s are show in S4 Fig). We calculated ΔAUC curves for each bacterial strain under increasing concentrations of each s100 (**Fig 3B**). Unlike the effect of human calprotectin, none of the four zebrafish s100 proteins exhibited nutritional immunity (Fig 3B; S4 Fig). Human calprotectin was the only protein to exhibit nutritional immunity under these conditions (Fig 3B, yellow curve).

Bacteria treated with zebrafish s100a1 showed improved growth relative to bacteria in the absence of s100 (Fig 3B, brown curve). Zebrafish s100a10b increased growth of *S. epidermidis* and had no effect on growth of zebrafish-derived bacterial strains (Fig 3B, orange). Zebrafish s100w (purple) improved *S. epidermidis* growth, showed a possible slight inhibitory effect on *Aeromonas ZOR001* for concentrations at or above 50 μM, and did not affect *Vibrio ZWU0020* growth at any concentration. Similarly, zebrafish icn (blue) improved *S. epidermidis* growth. We performed an ANOVA analysis on the antimicrobial activity at 50 μM protein for every condition and found that human calprotectin had a statistically different effect than any of the zebrafish s100 proteins (p < 0.0039), but that the zebrafish proteins could not be distinguished from one another (S3 Table).

### Zebrafish s100s do not exhibit proinflammatory activity like S100A9

Antimicrobial activity is not the only function of mammalian calprotectin. Calprotectin is a heterodimer of S100A8 and S100A9. The homodimer of mammalian S100A9 can potently activate an innate immune response through the Toll-like receptor 4 complex (TLR4), inducing nuclear localization of NF-κB and transcription of a wide variety of pro-inflammatory proteins (Foell et al. 2007; Björk et al. 2009). This activity can be reproduced in an *in vitro* functional assay by transfecting human embryonic kidney-293T (HEK293T) cells with plasmids encoding the proteins of the TLR4 complex (TLR4, MD-2, and CD14), as well as a plasmid placing luciferase behind an NF-κB promoter. We treat the cells with exogenous S100A9 and measure the expression and activity of luciferase produced in response (Loes, Bridgham, and Harms 2018).

We wanted to see if our purified zebrafish s100 proteins could play a similar role; therefore, we tested the ability of these proteins to activate TLR4 in this assay (**Fig 4**). Zebrafish have three ohnologs of tetrapod TLR4: Tlr4ba, Tlr4bb, and Tlr4al (Loes et al. 2021). Zebrafish Tlr4ba has been shown to induce inflammation in response to endotoxin, the small molecule lipopolysaccharide (LPS) derived from Gram-negative bacterial outer membranes, but neither Tlr4bb nor Tlr4al showed activity (Loes et al. 2021). We validated our assay by testing the ability of each complex to activate in response to endotoxin, the canonical agonist for the receptor (green). As expected, the human TLR4 and zebrafish Tlr4ba complexes responded strongly to endotoxin. As observed previously (Loes et al. 2021), Tlr4bb and Tlr4al did not show signal above vehicle treatment (light blue). We next challenged all four complexes with human S100A9 (yellow). We found that 2 μM human S100A9 activated human TLR4, as expected, but that none of the zebrafish Tlr4 complexes showed signal above background. This suggests that sterile inflammation via a mechanism similar to S100A9 activation of TLR4 is not a conserved function in zebrafish. Finally, we tested the ability of zebrafish s100s to activate each TLR4 complex at 2 μM. We observed no statistically significant agonist activity for any protein against any zebrafish Tlr4. We detected a small amount of signal for treatment with zebrafish s100a10b (orange) against human TLR4 (Bonferroni-corrected, paired two-sample t test of protein versus buffer, p value: 0.02); however, this was barely above background and much lower than the response for either endotoxin or human S100A9. We repeated the same experiment with 10 μM zebrafish s100s and observed no convincing agonist activity (S5 Fig).

These zebrafish s100 proteins show no evidence of activity against human TLR4 nor the three zebrafish Tlr4 ohnologs. Given the potent response of these receptors to positive controls (endotoxin or human S100A9), this strongly suggests proinflammatory activity is not present in the zebrafish s100 proteins tested. Further, with this assay, false positives are common due to endotoxin contamination from the recombinant expression of proteins in *E. coli*. The lack of signal thus gives strong evidence that these zebrafish proteins cannot activate TLR4 in the same fashion as human S100A9.

**Fig 4:**
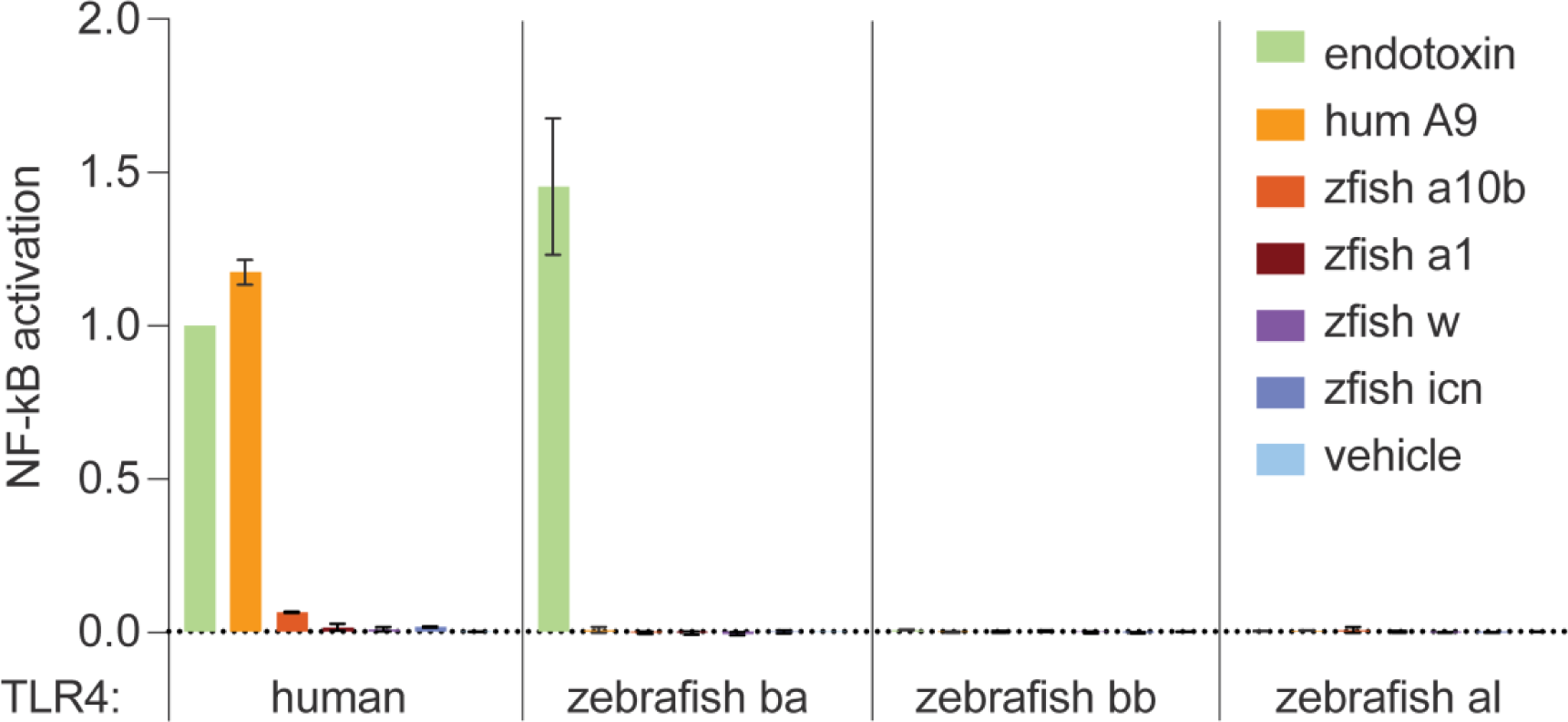
Zebrafish s100s do not exhibit the pro-inflammatory characteristics of S100A9. Activation of human and zebrafish TLR4 complexes in the presence of 2 μM zebrafish s100 proteins. Bars show the average signal across three biological replicates, with error bars indicating standard error. The positive controls for this experiment included human TLR4 and zebrafish Tlr4ba treated with endotoxin (green), and human TLR4 treated with 2 μM human S100A9 (yellow). There is no known agonist for zebrafish Tlr4bb and al complexes. All data was background subtracted and normalized to the signal from human TLR4 treated with endotoxin.

## DISCUSSION

Employing zebrafish as a model for studies of innate immunity and host-microbe interactions is a promising field of work. As vertebrates, zebrafish share much of their physiology and immune defense mechanisms with humans (Van Der Vaart, Spaink, and Meijer 2012), thus enabling mechanistic insight into health, disease, and the host-microbe interface. However, when approaching this research, we must be cognizant of the more than 400 million years since our most recent common ancestor, allowing for species-specific differences evolved to cope with diverse environments and other pressures.

We assert here that recent studies and commercial products have made the incorrect assumption that calprotectin exists in the fish innate immune response. We re-evaluate phylogenetic evidence to look for a homolog of calprotectin in fish and confirm that zebrafish do not share an s100 protein within the clade containing mammalian calprotectin.

We also tested the possibility that a fish s100 protein from a similar genomic context to calprotectin might have evolved calprotectin-like innate immune functions. We characterized the nutritional immunity and proinflammatory activity of four zebrafish s100s, including the previously studied zebrafish “calprotectin” s100a10b, in assays which are normally used to test calprotectin function. None of the zebrafish proteins performed like human calprotectin.

Although s100a10b is not calprotectin, researchers found that transcription of s100a10b increases in response to infection. This increase in expression is also reflected in the zebrafish injury scRNAseq dataset. But rather than viewing this as a convergent calprotectin or S100A8 activity, it seems best to view this through the lens of the mammalian S100A10 inflammatory response. This is because s100a10b is orthologous to S100A10; any shared activity between the human and zebrafish proteins likely reflects shared ancestry. Like zebrafish s100a10b, mammalian S100A10 is upregulated during inflammatory responses. While the function of zebrafish s100a10b is unknown, in mammals S100A10 regulates plasminogen-dependent macrophage migration via interactions with annexin A2. Based on our analyses, we believe such a role is a much more likely role for s100a10b than a convergently evolved calprotectin activity.

We cannot prove a negative: this work does not demonstrate that no zebrafish s100 protein exists that supplies some subset of the functions of human calprotectin. Our work does, however, put the burden of proof on researchers who would claim such functions exist. If a zebrafish s100 has antimicrobial or pro-inflammatory activity, it must have evolved that activity convergently and independently of those activities from mammalian calprotectin. Further, and importantly, such a protein does not shed direct light on mammalian biology. A convergent zebrafish s100 calprotectin-like protein would help us understand zebrafish biology and would be intriguing from the perspective of protein evolution. But it would almost certainly have a different regulatory scheme and subset of the human calprotectin functions.

We also did not test all possible zebrafish s100 proteins because we focused on those that seemed most promising to be expressed in immune cells during the immune response and shared genomic origin. Zebrafish s100 proteins represent a largely uncharacterized set of proteins, many of which evolved in the ray-finned fishes (Kraemer, Saraiva, and Korsching 2008). We are excited to see how investigations of their functional roles continue. We remain intrigued by the idea that convergent evolution may exist between mammalian calprotectin and some other protein(s) in zebrafish.

In addition to studying zebrafish s100 proteins individually, one important line of work will be to investigate various s100 heterocomplexes. Mammalian calprotectin is a heterodimer and heterotetramer, but we only performed experiments with homodimeric zebrafish s100s. Studies of other heterodimeric human S100 complexes have been done and prove to have altered functions (Spratt et al. 2019). In the future, this type of analysis could be done with zebrafish proteins to explore whether a heterodimer state confers nutritional immunity or proinflammatory activity. However, there is currently no evidence at this point suggesting this is likely.

We conclude that it is crucial that we use an evolutionary lens and careful biochemical analyses to probe homology between zebrafish and human proteins so that we can make accurate extrapolations of findings from zebrafish models of human biology.

## MATERIALS AND METHODS

### Sequence analysis

In our BLAST and reciprocal BLAST study, we used the NCBI web interface and BLASTP with default parameters to query the NCBI RefSeq database (v. 244). We aligned and calculated S100 sequence identity using Clustal Omega (Madeira et al. 2022). We viewed and generated figures displaying the alignment and consensus sequence in SnapGene Viewer 5.2.4.

### Protein Purification

We purchased all zebrafish s100 genes from GenScript in the pET-28a(+)-TEV vector with an N-terminal 6x-Histidine tag and TEV protease cleavage site (S2 Table). All genes were codon-optimized for expression in *E. coli*. We expressed human calprotectin (with S100A9 containing the C3S mutation) and human S100A9/C3S in a pET-Duet vector without purification tags. We transformed Rosetta2(DE3)pLysS *E. coli* cells with plasmids. We used transformant glycerol stocks to inoculate cultures in 15 mL Luria broth (LB) with 50 μg/mL kanamycin and 34 μg/mL chloramphenicol. We incubated cultures overnight at 37 °C, shaking at 250 rpm. The following day, we diluted 15 mL saturated cultures into 1.5 L of LB with antibiotics. When the OD_600_ reached 0.6-1.0, we induced recombinant protein expression with 1 mM IPTG and 0.2% glucose and then grew overnight at 16 °C, shaking at 250 rpm. We pelleted cells at 3,000 rpm for at least 15 minutes in an F6B rotor in a Beckman Coulter preparative centrifuge. We stored pellets at -20 °C for up to one month.

We prepared protein lysates for purification with the following method: We vortexed pellets (6-9 g) in 45 mL buffer from the first chromatography step (see below) until cells were resuspended, added 15 µL each of DNase I and Lysozyme (ThermoFisher Scientific), and incubated at room temperature with gentle shaking for at least 10 minutes. We lysed the cells by sonication at 55% amplitude with 0.3 second pulse on, 0.7 second pulse off, for 3-5 minutes. We pelleted cell debris by centrifugation at 15,000 rpm at 4 °C for at least 20 minutes in a JA-20 rotor in a Beckman Coulter preparative centrifuge and collected the supernatant. To remove remaining large debris, we filtered lysate supernatant through a 0.2 µm pore syringe filter immediately prior to purification chromatography.

We purified all proteins using an Äkta PrimePlus Fast Protein Liquid Chromatography system using two stacked 5 mL HiTrap columns at each step. We used HisTrap FF columns for Ni-affinity and Q HP columns for anion exchange (GE Health Science). All chromatography was performed at 4 °C. At the end of purification, we confirmed protein purity was >95% by SDS-PAGE. Then, we dialyzed each protein overnight into 4 L of 25 mM Tris, 100 mM NaCl, pH 7.4 at 4 °C. We placed 2 g/L Chelex 100 resin (Bio-Rad) in the dialysis buffer to chelate divalent metal ions. We concentrated each protein to roughly 2 mg/mL and syringe-filtered through a 0.22 µm filter directly into liquid nitrogen to sterilize and flash freeze before storing at -80 °C.

We purified TEV-cleavable 6xHis-tagged zebrafish s100 proteins with the following scheme. We used 25 mM Tris, 100 mM NaCl, pH 7.4 buffer as the base for all chromatography buffers. We ran our protein lysate over a Ni-affinity column with a 50 mL wash and eluted over a 75 mL gradient from 25-1000 mM imidazole to collect proteins with strong Ni binding capacity. We determined which fractions contained our desired protein by SDS-PAGE and pooled these fractions. To separate our recombinant proteins from their Ni-binding His-tag, we added 5 mM DTT and 6xHis-tagged TEV protease to the pooled fractions and incubated the reaction at room temperature with gentle shaking for at least 5 hours. We then dialyzed the protein solution overnight into 4 L buffer with 25 mM imidazole and 5 mM DTT to allow cleavage to come to completion and to remove excess imidazole from the sample. We performed a second round of Ni-affinity chromatography. Without the His-tag, the zebrafish s100s have low affinity for Ni. Therefore, we isolated pure, non-tagged zebrafish s100 proteins at this step during a 50 mL wash in 25 mM imidazole and then used a step gradient to 1 M imidazole to elute His-tagged and other contaminant proteins that had higher affinity for the Ni column. Purified zebrafish s100s were prepared for storage as described above.

We purified human calprotectin using Ni-affinity chromatography at pH 7.4 and anion exchange at pH 8. When expressing calprotectin, S100A8 and S100A9 homodimers are also expressed and must be removed during chromatography. In the presence of calcium, S100A9 and calprotectin bind divalent metal ions like Ni, but S100A8 and most other lysate proteins do not. We loaded our Ni-affinity column with calprotectin lysate, washed away most A8 and contaminants in a 50 mL wash, and then eluted calprotectin and S100A9 over a 75 mL gradient from 0-1000 mM imidazole and 1-0 mM CaCl_2_ in 25 mM Tris, 100 mM NaCl, pH 7.4 buffer.

We pooled elution peak fractions containing calprotectin and contaminant S100A9 homodimers, as determined by SDS-PAGE, and dialyzed overnight in 4 L of 25 mM Tris, 100 mM NaCl at pH 8. We loaded our sample onto an anion exchange chromatography column in 25 mM Tris, 100 mM NaCl at pH 8 with 100 mM NaCl. Because S100A9 has a lower pI than calprotectin, it binds the anion column more strongly at pH 8. We used a 50 mL wash in 100 mM NaCl to isolate calprotectin and then used a step gradient increasing the salt to 1 M NaCl to remove S100A9 and other contaminants from the column. At this point, calprotectin was pure and prepared for storage as described, and fractions with S100A9 and other contaminants were discarded.

We used a similar protocol to purify human S100A9. We performed Ni-affinity chromatography as described for calprotectin. We then performed anion exchange chromatography using a 50 mL wash and collected fractions over a 70 mL gradient elution from 100-1000 mM NaCl in 25 mM Tris, pH 8 buffer to isolate S100A9 from contaminant proteins that also bind the anion column. We used SDS-PAGE to confirm fractions with S100A9, pooled and dialyzed these fractions overnight into 4 L of 25 mM Tris, 100 mM NaCl at pH 6. As a final step, we loaded the S100A9 sample onto an anion exchange column in 25 mM Tris, 100 mM NaCl buffer at pH 6. S100A9 binds weakly to the anion column at pH 6. Therefore, we collected S100A9 in a 50 mL wash in 100 mM NaCl, and then removed contaminants from the column with a step elution at 1 M NaCl. Pure S100A9 was prepared for storage as described.

### Far-UV Circular Dichroism and Fluorescence Spectroscopy

Prior to biophysical measurements, we thawed and equilibrated all proteins into 25 mM Tris, 100 mM NaCl, pH 7.4 via overnight dialysis in 4 L buffer at 4 °C. We determined protein concentrations by Bradford Assay using bovine serum albumin (BSA) standards and the molecular weight of each dimeric structure, then diluted to ∼10 µM in dialysis buffer.

For all spectroscopic measurements, we assessed metal-induced changes to the spectra by measuring the spectrum in the presence of 2 mM CaCl_2_ and then adding excess EDTA at 5 mM and re-measuring the spectrum. We collected far-UV circular dichroism data between 200–250 nm using a J-815 CD spectrometer (Jasco) with a 1 mm quartz spectrophotometer cell (Starna Cells, Inc. Catalog No. 1-Q-1). We collected 3 scans for each condition, and then averaged the spectra and subtracted a blank buffer spectrum using the Jasco spectra analysis software suite. We converted raw ellipticity into mean molar ellipticity using the concentration and number of residues in each protein.

We collected intrinsic tyrosine and/or tryptophan fluorescence using the J-815 CD spectrometer (Jasco) with an attached model FDT-455 fluorescence detector (Jasco) using a 1 cm quartz cuvette (Starna Cells, Inc.). We collected a single excitation and emission scan at 10 nm/min with a 10 nm bandwidth, 1 nm data pitch, and 1 sec D.I.T. for each condition and then subtracted a blank buffer spectrum using the Jasco spectra analysis software suite. Depending on the sample signal, we set the detector sensitivity to either 630 or 800 Volts. We conducted excitation scans by measuring 305 nm light emitted at for all zebrafish proteins and 345 nm emitted light for human S100A9 for each excitation wavelength from 200-295 nm. For emission scans we used 280 nm light to excite zebrafish proteins and 288 nm for human S100A9, and measured light emitted at all wavelengths from 285-425 nm.

### Nutritional Immunity Assay

We measured the antimicrobial activity of zebrafish s100s and human calprotectin against human- and zebrafish-derived bacterial strains using a modified version of a well-established assay that will be described here (Brophy, Hayden, and Nolan 2012; Nakashige et al. 2016; Hadley et al. 2018; Harman et al. 2020). Bacterial strains used in this assay include 1) *Staphylococcus epidermidis,* a human commensal strain previously shown to respond to calprotectin (Hadley et al. 2018; Harman et al. 2020); 2) *Aeromonas ZOR001*, isolated from zebrafish and not previously characterized for response to calprotectin; and 3) *Vibrio ZWU0020*, isolated from zebrafish and not previously characterized for response to calprotectin but related to human-derived *Vibrio cholerae* shown to respond to calprotectin (Farr et al. 2022). We obtained both zebrafish-derived strains from the Guillemin lab at the University of Oregon.

Each week, we plated bacterial strains from glycerol stocks onto antibiotic-free LB agar and grew at 30 °C overnight before storing plates at 4 °C. The day before an experiment, we inoculated a 5 mL culture in liquid LB media with a single colony from each strain and grew overnight at 30 °C with shaking. The following day, we diluted cultures 1:100 in 5 mL LB and grew to an OD_600_ around 0.8 by the time of the experiment. *Aeromonas ZOR001* and *Vibrio ZWU0020* were diluted 2 hours before the experiment. *S. epidermidis* grew more slowly so required dilution 4 hours prior to the experiment.

The day before each experiment, we thawed a single S100 protein from -80 °C, concentrated to at least 200 μM using a Nanosep 3K Omega spin concentrator (Pall Corporation), and dialyzed overnight at 4 °C into 4 L of Experimental Buffer (25 mM Tris, 100 mM NaCl, pH 7.4) with 2 g/L Chelex 100 resin (Bio-Rad) to chelate residual transition metal ions. After dialysis, we filter-sterilized the protein through a Ultrafree-MC-VV centrifugal filter with Durapore PVDF 0.1 µm and kept at 4 °C until time of experiment.

To start the experiment, we made a protein dilution series by mixing a desired amount of protein in sterile Experimental Buffer with the appropriate amount of LB to achieve a ratio of 62:38, respectively. We then brought the volume of these protein solutions up to 1.7 mL in Experimental Media (EM). We made EM by mixing Experimental Buffer in LB at a ratio of 62:38, respectively, and filter-sterilized. We distributed each sample in aliquots of 160 µL across ten wells of a clear Falcon 96-Well, Cell Culture-Treated, Flat-Bottom Microplate. At this time, we diluted each bacterial strain to an estimated OD_600_ of .008 in 5 mL Experimental Media with calcium (EMC). We made EMC by adding 10.2 μM CaCl_2_ to EM, and sterile-filtered. Then, we added 40 µL of dilute bacteria or EMC without bacteria (contamination control) to each well, bringing the final volume to 200 µL/well, and making technical triplicate conditions for bacterial strains. To counteract sample evaporation, the outermost wells of the plate contained 160 μL EM and 40 μL EMC, and we wrapped the plate in a single layer of parafilm.

We measured bacterial growth by OD_600_ every 15 minutes over 13 hours in a Molecular Devices SpectraMax i3. The plate was shaken for 5 seconds before the first read, then for 10 minutes between each subsequent read. We set the plate reader temperature to 25 °C, however, over the course of the overnight growth, the actual temperature reached 37 °C. The final concentration of metals in the media without bacteria was measured using ICP-MS at the USR Elemental Analysis Core. The measured concentrations were Ni: 45.4 μM, Ca: 107.3 μM, Cu: 157.4 μM, Mg: 160.5 μM, Mn: 216.6 μM, Fe: 1.1 mM, and Zn: 5.9 mM.

For the analysis, we background subtracted each experimental condition using OD_600_ values for the matching concentration of S100 protein concentration in buffer without bacteria added. We used Prism to average the replicates by condition, determine the standard error of the mean, and graph the results.

### Proinflammatory Activity Assay

We tested the S100A9-like proinflammatory activity of zebrafish s100s using a well-established assay (Loes, Bridgham, and Harms 2018; Loes et al. 2021; Harman et al. 2020). This assay measures relative activation of the TLR4-mediated immune response through NF-κB. For each experiment, we thawed all zebrafish s100 proteins and human S100A9 from -80 °C, buffer exchanged into endotoxin-free PBS, then treated with endotoxin removal spin columns (ThermoFisher Scientific) to remove residual LPS from the purification process.

We performed each experiment in technical triplicate and followed the Dual-Glo Luciferase Assay System protocol (Promega). We transiently transfected adherent HEK293T cells (ATCC; CRL-11268) in a Falcon 96-Well, Cell Culture-Treated, Flat-Bottom Microplate with pcDNA vector plasmids using PLUS and Lipofectamine Reagents (ThermoFisher Scientific). Plasmids contained genes for human or zebrafish TLR4 complex components and *Renilla* luciferase enzyme under constitutively active promoters, and the firefly luciferase gene controlled by an NF-κB promoter. For human TLR4 complex transfections, we transfected 10 ng human TLR4, 0.5 ng human MD-2, and 1 ng human CD14 plasmids per well. For zebrafish Tlr4 complex transfections, we used 10 ng zebrafish Tlr4ba, bb, or al, 20 ng zebrafish Md-2, and 1 ng mouse CD14 plasmids per well, as this ratio gives us the best signal to noise ratio. Zebrafish do not have an annotated CD14. Previous studies have shown zebrafish Tlr4ba can be activated in the presence of mouse and human CD14, but more strongly with mouse. We also transfected all wells with 1 ng *Renilla* plasmid, 20 ng elam-Luc (firefly), and brought the total DNA mass per well to 100 ng with empty pcDNA vector in a total media volume of 200 μL per well.

After 20-24 hours incubation at 37 °C in 5% CO_2_, we removed all 200 µL of transfection mix from each well. We then treated transfected HEK293T cells with 100 µL of one of the following treatment mixes: 1) 2 µM S100 protein and 200 ng/μL Polymyxin B to bind up LPS in media, 2) 0.2 ng/µL LPS-R (tlrl-eklps; Invivogen) as a positive control for human TLR4, or 3) 2 ng/µL lipid IVa as a positive control for zebrafish Tlr4ba activation. Because there is no known activator of zebrafish Tlr4bb and Tlr4al complexes (Loes et al., 2021), we treated these transfected cells with 2 ng/µL lipid IVa for consistency. After incubating again at 37 °C in 5% CO_2_ for 3-4 hours, we removed and discarded 60 µL of treatment mix from each well. We chemically lysed the cells by adding 30 µL Dual Glo lysis reagent containing firefly luciferin and incubated in the dark for 7 minutes. We then mechanically lysed the cells by scraping the bottom of each well with a pipet tip and transferring 60 µL of cell solution to an opaque 96-well plate. After a 7-minute incubation in the dark at room temperature, we measured luminescence per well produced by firefly luciferase activity using a Molecular Devices SpectraMax i3. Then we added 30 μL of Dual-Glo Stop & Glo buffer containing firefly luciferase quencher and *Renilla* luciferase reagent, incubated for 7 more minutes, and measured luminescence.

For the analysis, we took the firefly signal for each experimental condition and background subtracted the averaged firefly signal of wells transfected with the corresponding complex but treated with buffer without agonist. We did the same for the *Renilla* signal, with background signal considered as the averaged signal from wells with same treatment condition but transfected only with vector. We divided the background-subtracted firefly signal for each well by the background-subtracted *Renilla* signal for that same well. To simplify comparisons across experiments, we normalized the firefly/*Renilla* value for each well to the triplicate average of the firefly/*Renilla* values for human TLR4 complex treated with 0.2 ng/μL LPS-R.

## ACKNOWLEDGMENTS

This work was funded by NIGMS R01-GM146114 (MJH). The funders had no role in study design, data collection and interpretation, or the decision to submit the work for publication. We would like the thank the Guillemin lab at the University of Oregon for the zebrafish-derived bacterial strains used in this work, as well as guidance on experiments and interpretation of data.

