## Supplemental Material for "Zebrafish do not have calprotectin"

**SUPPLEMENT**

| | A8 | A9 | A10 | a10b | w | icn | icn2 | a1 | t |
|---|---|---|---|---|---|---|---|---|---|
| A8 | 100 | 25.81 | 27.78 | 34.41 | 29.35 | 30.00 | 30.00 | 32.61 | 34.41 |
| A9 | | 100 | 29.90 | 37.00 | 29.13 | 33.68 | 36.08 | 39.58 | 33.33 |
| A10 | | | 100 | 52.58 | 27.08 | 38.46 | 36.26 | 47.31 | 42.70 |
| a10b | | | | 100 | 33.33 | 43.62 | 42.55 | 54.17 | 46.74 |
| w | | | | | 100 | 30.11 | 29.03 | 33.68 | 29.47 |
| icn | | | | | | 100 | 87.37 | 43.62 | 50.00 |
| icn2 | | | | | | | 100 | 87.37 | 51.16 |
| a1 | | | | | | | | 100 | 54.55 |
| t | | | | | | | | | 100 |

**S1 Table:** Identity matrix for human S100 proteins (A8, A9, and A10) and homologs from zebrafish (a10b, w, icn, icn2, a1, and t).

| Gene | Vector | Protein sequence |
|---|---|---|
| Human Calprotectin<br><br>NP_001306130.1<br>(S100A8/C42S) and<br>NP_002956.1<br>(S100A9/C3S) | pETDUET | MLTELEKALNSIIDVYHKYSLIKGNFHAVYRD<br>DLKKLLETE**S**PQYIRKKGADVWFKELDINTD<br>GAVNFQEFLILVIKMGVAAHKKSHEESHKE<br><br>MT**S**KMSQLERNIETIINTFHQYSVKLGHPDTL<br>NQGEFKELVRKDLQNFLKKENKNEKVIEHIM<br>EDLDTNADKQLSFEEFIMLMARLTWASHEKM<br>HEGDEGPGHHHKPGLGEGTP |
| Human S100A9<br>NP_002956.1/C3S | pETDUET | MT**S**KMSQLERNIETIINTFHQYSVKLGHPDTL<br>NQGEFKELVRKDLQNFLKKENKNEKVIEHIM<br>EDLDTNADKQLSFEEFIMLMARLTWASHEKM<br>HEGDEGPGHHHKPGLGEGTP |
| Zebrafish s100a10b<br>NP_998168.1 | pET-28a(+)-TEV | mgsshhhhhhssgenlyfqghMPSDLERAMETLITVFH<br>RYSGAEGNSSTLSRRELKQLMEKELASFLKSQ<br>KDPAAVDKIMKDLDANGDGEVNFEEFVSLVV<br>GLSIACEQLYQKQMKAAAKK |
| Zebrafish s100a1<br>NP_001082820.1 | pET-28a(+)-TEV | mgsshhhhhhssgenlyfqghMVSKLENAMEGLIKVF<br>HTYSSKEGDKYKLSKSELKSLLQGELSDFLAA<br>SKDPMVVEKIMSDLDENRDGEVDFQEFVVLV<br>AALTVACNEFFVESMKN |
| Zebrafish s100w<br>NP_001313380.1 | pET-28a(+)-TEV | mgsshhhhhhssgenlyfqghMSKLEKAIVSIVEVFEE<br>YAGKDEQKSQLSNAELRDLIKTELKSPEFKDK<br>VDPENIKEVMEELDKNHDGEVNFREFSQCIAG<br>LARGYYKKKHGNEERRGRGRDK |
| Zebrafish icn<br>NP_997926.1 | pET-28a(+)-TEV | mgsshhhhhhssgenlyfqghMATSDTQKAMAMLIAT<br>FHKYSGKEGDKLTLSKGELKELLSAELGDIFG<br>KTTDKAALDKIFKDLDANADGSVDFQEYITLI<br>ACITMLCNEFFTGKK |

**S2 Table:** Sequences of s100 genes expressed and purified in this study. Lowercase amino acids indicate the addition of a 6xHis tag and TEV protease cleavage site. Bold and underlined amino acids are mutations relative to the reference sequence.

| level | degrees of freedom | sum of squares | mean square | F-value | p-value |
|---|---|---|---|---|---|
| **species** | **2** | **34.6** | **17.3** | **11.7** | **0.004** |
| **protein** | **4** | **80.1** | **20.0** | **13.6** | **0.001** |
| Residuals | 8 | 11.8 | 1.5 | --- | --- |

| level | comparison | difference | lower | upper | p-value |
|---|---|---|---|---|---|
| **species** | ***S. epi.* vs. *A. ZOR001*** | **3.72** | **1.52** | **5.91** | **3.25E-03** |
| species | *V. ZWU0020* vs. *A. ZOR001* | 1.75 | -0.45 | 3.94 | 1.18E-01 |
| species | *V. ZWU0020* vs. *S. epi.* | -1.97 | -4.16 | 0.22 | 7.66E-02 |
| **protein** | **zf s100a1 vs. hCP** | **6.17** | **2.75** | **9.59** | **1.69E-03** |
| **protein** | **zf s100a10b vs. hCP** | **5.73** | **2.31** | **9.15** | **2.74E-03** |
| **protein** | **zf ictacalcin vs. hCP** | **5.42** | **1.99** | **8.84** | **3.93E-03** |
| **protein** | **zf s100w vs. hCP** | **5.67** | **2.24** | **9.09** | **2.94E-03** |
| protein | zf s100a10b vs. zf s100a1 | -0.44 | -3.87 | 2.98 | 9.90E-01 |
| protein | zf ictacalcin vs. zf s100a1 | -0.76 | -4.18 | 2.67 | 9.34E-01 |
| protein | zf s100w vs. zf s100a1 | -0.50 | -3.93 | 2.92 | 9.84E-01 |
| protein | zf ictacalcin vs. zf s100a10b | -0.31 | -3.74 | 3.11 | 9.97E-01 |
| protein | zf s100w vs. zf s100a10b | -0.06 | -3.48 | 3.36 | 1.00E+00 |
| protein | zf s100w vs. zf ictacalcin | 0.25 | -3.17 | 3.68 | 9.99E-01 |

**S3 Table: Results of an ANOVA analysis of zebrafish s100 protein antimicrobial activity.** The top sub-table shows the correlation of bacterial species (*S. epi.*, *A. ZOR001*, and *V. ZWU0020*) and protein (hCP, s100a1, s100a10b, ictacalcin, s100w) with the mean change in area under the growth curves for the 50 μM treatment condition. The bottom sub-table shows the results of a post hoc Tukey test applied to the ANOVA results. This test reveals that the effect of hCP is significantly different than any of the zebrafish proteins (p values between 0.00169 and 0.0039; bolded rows). The effects of the zebrafish proteins cannot be distinguished.

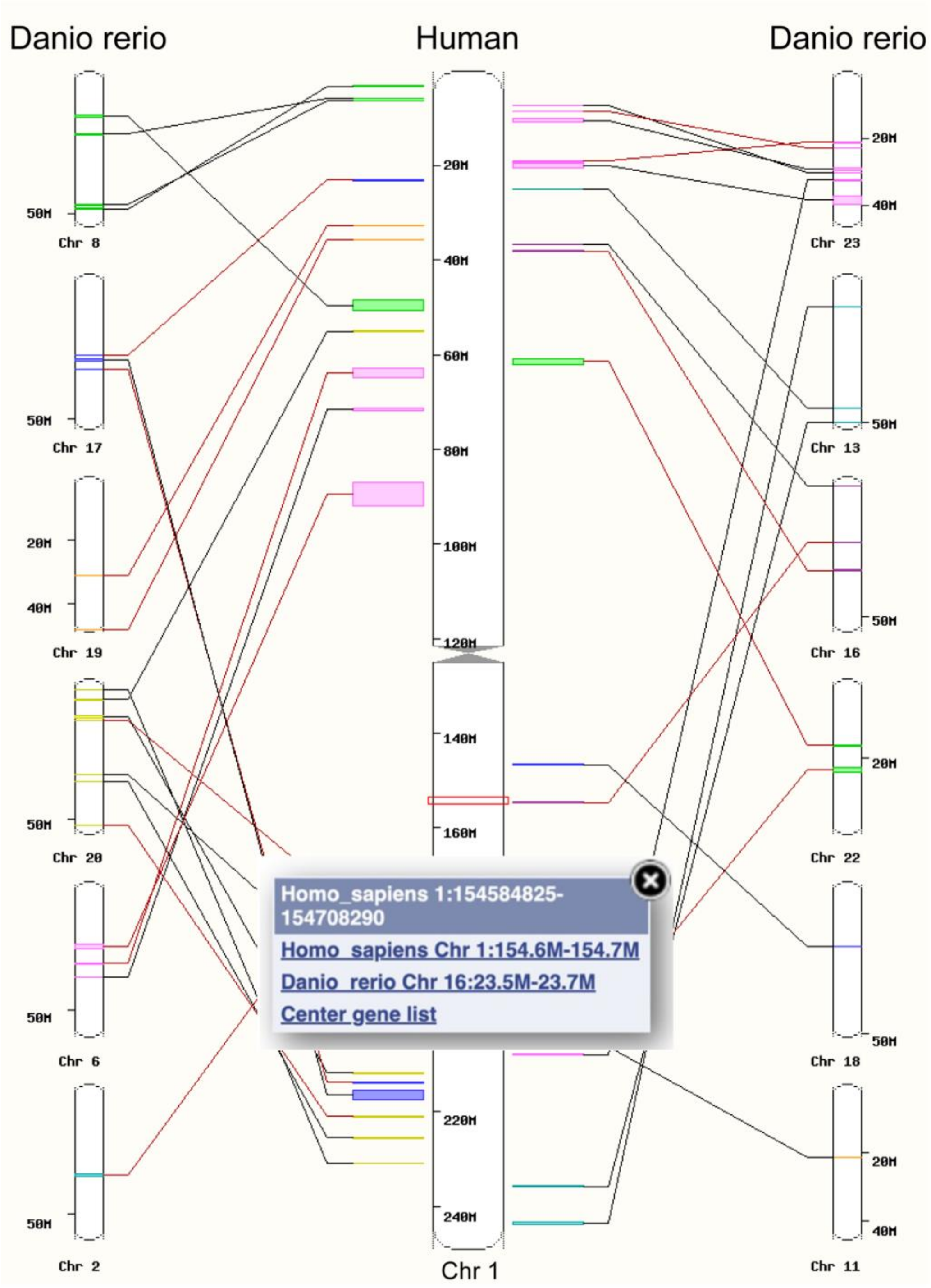


**S1 Fig: EMSEMBL identification of synteny between region of human S100 genes and the zebrafish genome**. Central chromosome is human chromosome 1; outer chromosomes are zebrafish chromosomes with regions syntenic to regions of human Chr1. The red box on Chr1 indicates the region containing 19 of the 24 human S100 genes (1:154584825-154708290). This is syntenic to zebrafish Chr16.

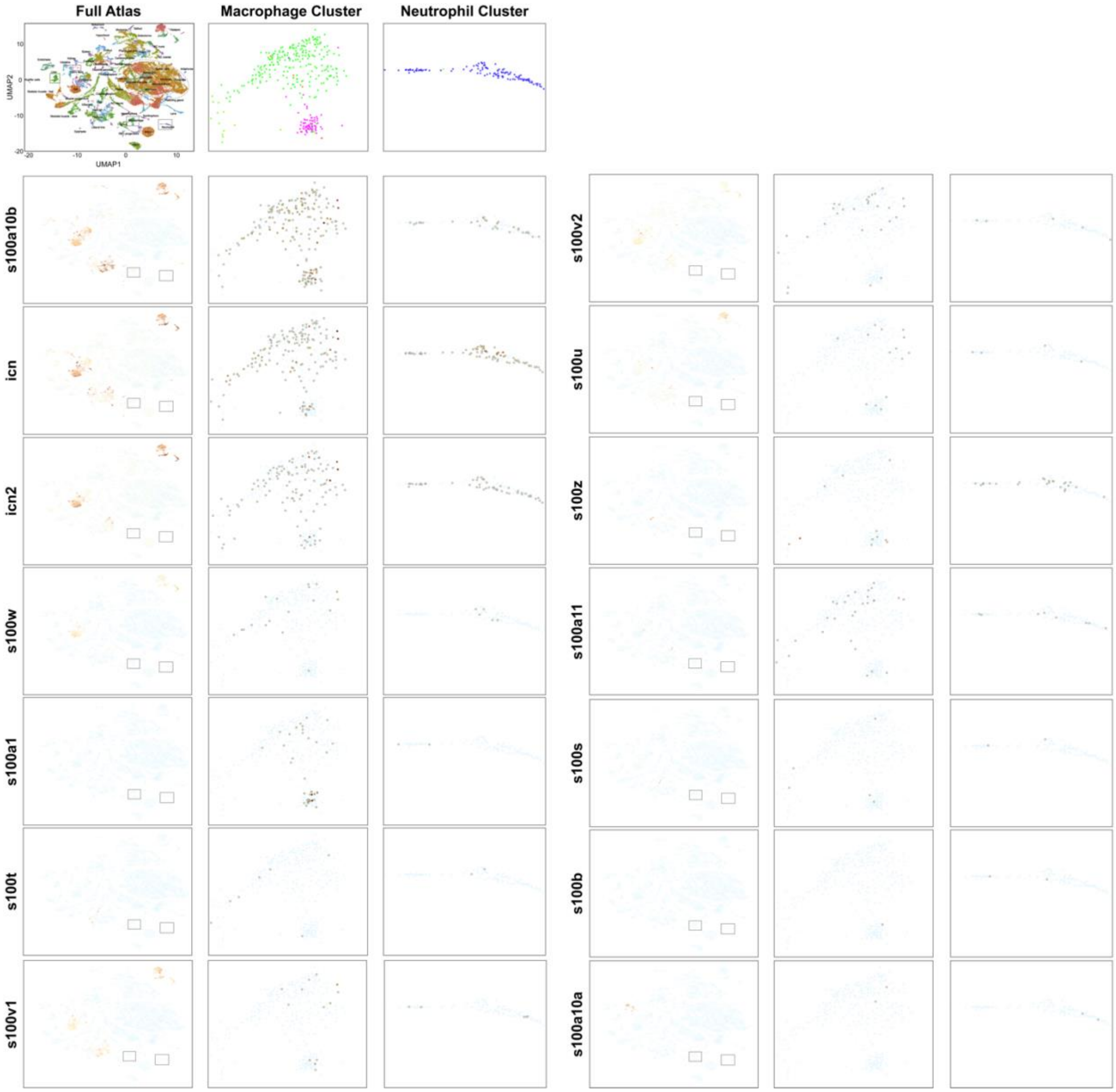


**S2 Fig: Single-cell transcription of zebrafish s100 genes in immune cells.** Developmental scRNAseq dataset mining taken from the UCSC cell browser NCBI Bioproject: 564810 (Farnsworth, Saunders, and Miller 2020; Speir et al. 2021). Within each panel, each point is a cell, separated by transcriptional profile along the UMAP1 and UMAP2 axes. The left-most panels show all cells; the middle- and right-most panels zoom in on the regions corresponding to macrophages and neutrophils. These regions are shown by the small boxes on the left-most panels. Each row of panels corresponds to a different zebrafish s100 gene (labeled on the left). Cells are colored by the relative expression level of that gene. Rows are sorted from highest expression to lowest.

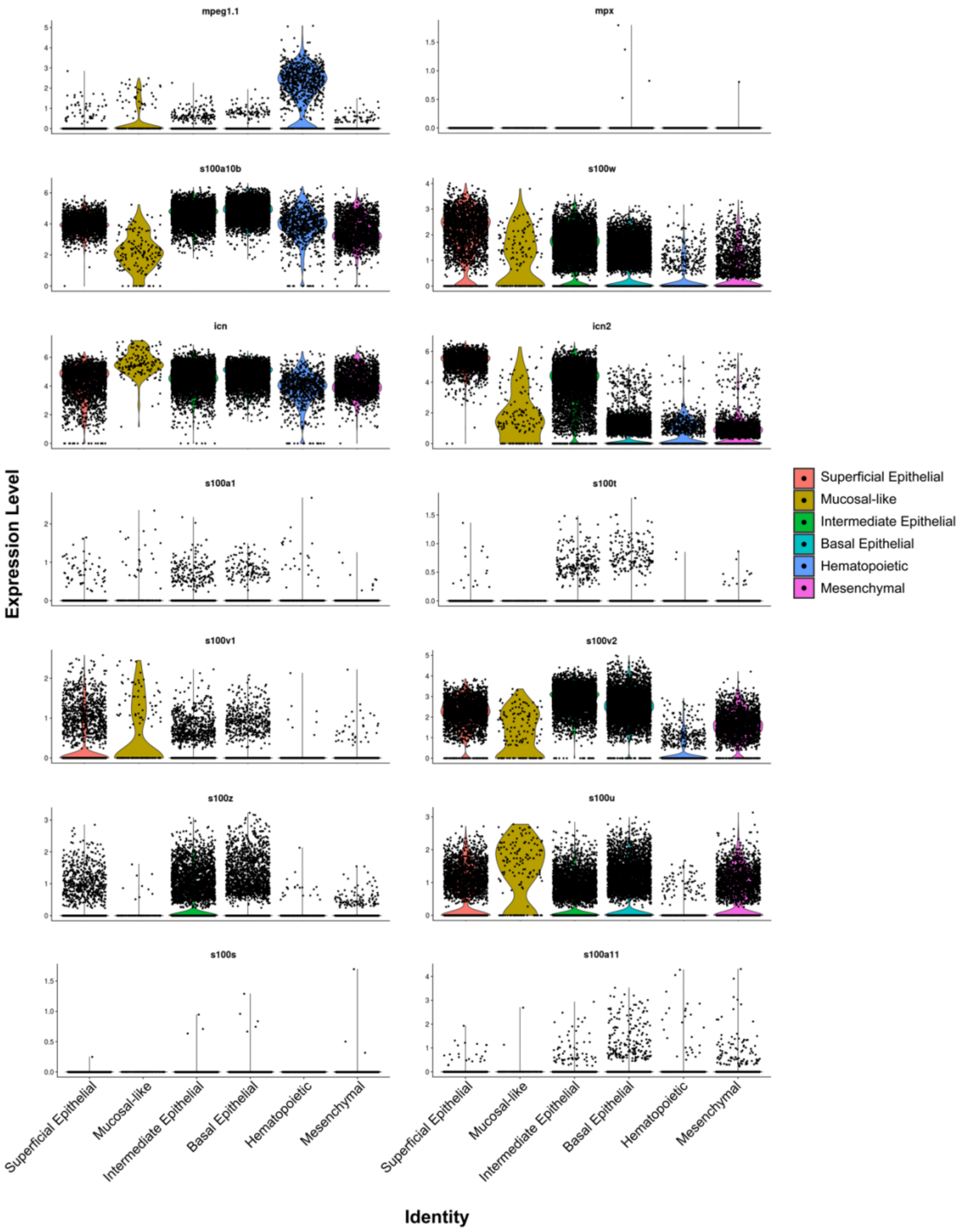


**S3 Fig. Single-cell transcription of zebrafish s100 genes in response to injury (fin clip).** Data are taken from NCBI GEO accession number GSE137971 (Hou et al. 2020). We accessed the data they published from *https://k326xh.shinyapps.io/FinRegenerationSCRNA/* on June 11$^{th}$, 2024. Panels show the change in the transcription of specific genes in response to fin clip. Panel title indicates gene in question (e.g., mpeg1.1, mpx, s100a10b, etc.). Within each panel, series show response in specific tissue types, labeled at the bottom. The y-axis is the gene expression level. Individual cells are shown as points.

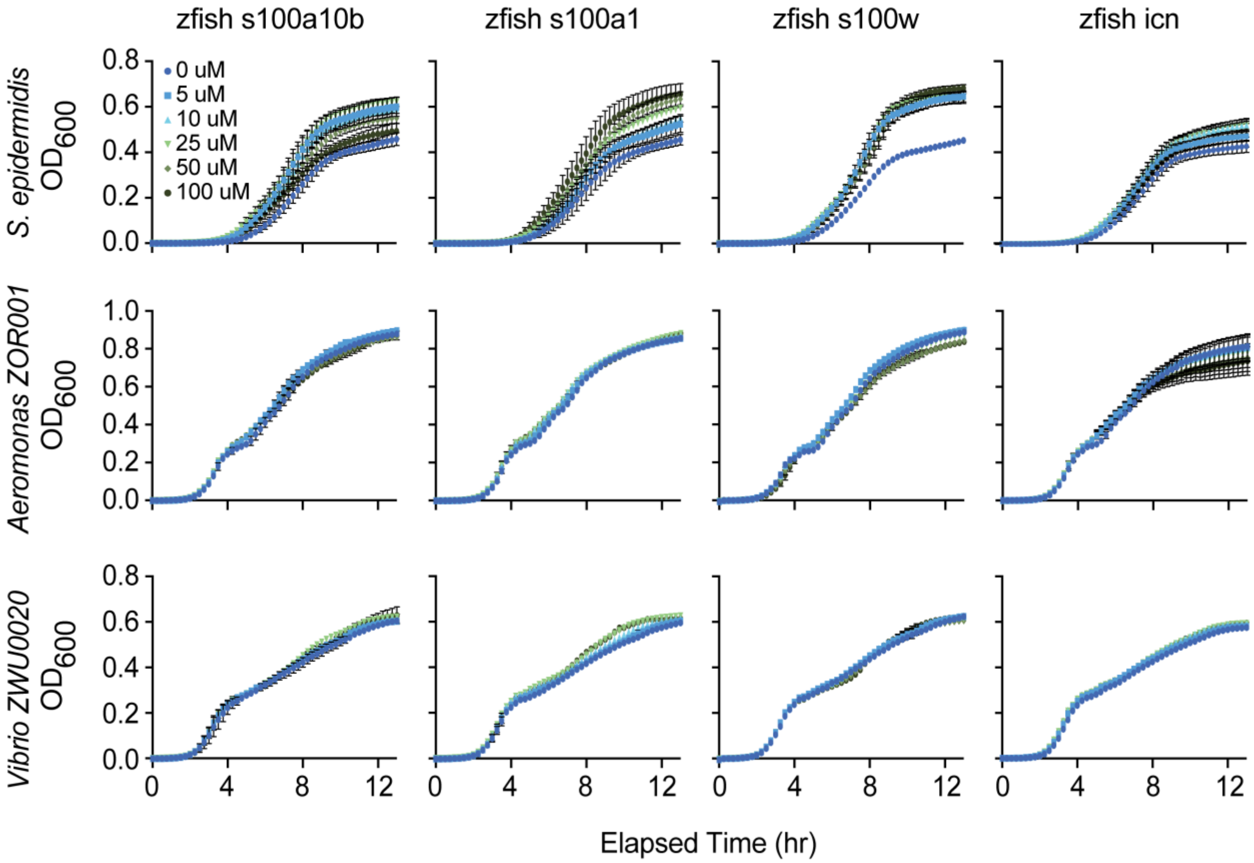


**S4 Fig. Nutritional immunity assays of zebrafish s100s.** Rows show assays done with the bacterial strain indicated on the left. Columns show results for the S100 protein indicated at the top. Bacterial growth was measured over 13 hours in the presence of S100 concentrations ranging from 0-100 µM, dark blue to dark green as shown in the legend at the top left. All measurements were done in biological triplicate of technical triplicates except s100A1 at 100 µM which only contains data from two biological replicates. Datapoints and error bars represent the mean and standard error of biological replicates.

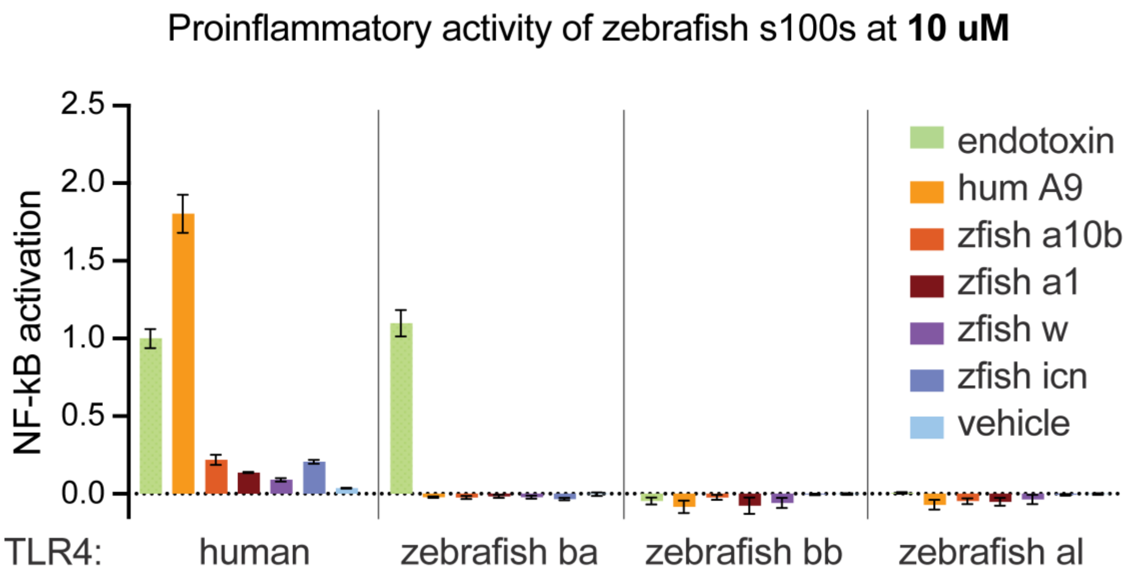


**S5 Fig. Proinflammatory assay with zebrafish s100 concentrations increased 5-fold.** Bars show the average signal across three biological replicates, with error bars indicating standard error. The positive controls for this experiment included human TLR4 and zebrafish TLR4ba treated with endotoxin (green), and human TLR4 treated with 2 μM human S100A9 (yellow). For zebrafish experiments, we used 10 μM protein. There is no known agonist for zebrafish TLR4bb and al complexes. All data was background subtracted and normalized to the signal from human TLR4 treated with endotoxin.
